# Cell cycle control of cohesion establishment

**DOI:** 10.1101/2025.04.18.649605

**Authors:** Brooke D. Rankin, Vanessa A. Moore, Damira Flavors, Susannah Rankin

**Affiliations:** Cell Cycle and Cancer Biology Program, Oklahoma Medical Research Foundation, Oklahoma City, OK 73104, USA; Cell Biology Department, University of Oklahoma Health Sciences Center, Oklahoma City, OK 73114

## Abstract

Cohesion between sister chromatids ensures accurate alignment and segregation of chromosomes during cell division. Cohesion establishment requires modification of the SMC3 subunit of cohesin by the ESCO2 acetyltransferase. ESCO2 function relies on interactions between ESCO2 and the DNA replication machinery, but how and whether these interactions are regulated during cell cycle progression is not known. Here we have found that phosphorylation of ESCO2 at serine 75 by cyclin-dependent kinase strongly impacts both interaction of ESCO2 with the replication machinery *and* ESCO2’s ability to ensure sister chromatid cohesion. This study provides mechanistic insight into how establishment of sister chromatid cohesion is entrained in S phase by cell cycle-dependent mechanisms.

## Introduction

During eukaryotic DNA replication chromosomes are duplicated to form two identical copies called sister chromatids, which remain tethered together until they are pulled apart at anaphase. This tethering, called sister chromatid cohesion, ensures satisfaction of the spindle checkpoint and progression through mitosis, and thus promotes proper chromosome segregation (Tanaka et al., 2000; Uhlmann et al., 1999; Hoque and Ishikawa, 2002). Additionally, sister chromatid cohesion plays a critical role in certain kinds of DNA repair based on homologous recombination, thus ensuring maintenance of genomic integrity (Sjögren and Nasmyth, 2001; Ström et al., 2004).

Sister chromatid cohesion is mediated by cohesin, a multi-subunit protein complex that associates with chromatin throughout most of the cell cycle (Sumara et al., 2000). Cohesin is loaded onto chromosomes in telophase, and this association remains dynamic throughout G1 (Gerlich et al., 2006; Kueng et al., 2006). In vertebrates, the establishment of sister chromatid cohesion during DNA replication requires the acetyltransferase ESCO2 (Higashi et al., 2012; Song et al., 2012; Lafont et al., 2010; Hou and Zou, 2005; Kawasumi et al., 2017). This enzyme modifies the SMC3 subunit of cohesin, rendering the cohesin complex resistant to removal from chromatin, in a mechanism conserved with lower eukaryotes (Zhang et al., 2008; Ünal et al., 2008; Rowland et al., 2009; Ben-Shahar et al., 2008). Although cohesin stabilization can also be promoted by the activity of a related enzyme called ESCO1, in previous work we showed that sister chromatid cohesion is critically dependent on ESCO2 (Alomer et al., 2017).

ESCO2 protein levels are controlled at least in part in a cell cycle-dependent manner. ESCO2 is targeted for degradation in G1 by the anaphase-promoting complex (APC), and ESCO2 levels can be seen to rise during S phase when the APC is inactive (Lafont et al., 2010; Song et al., 2012; Minamino et al., 2018). Some studies suggest that ESCO2 may be a target of Cul4-dependent degradation in late S, although alternative models have also been proposed (Jevitt et al., 2023; Minamino et al., 2018).

The ability of ESCO2 to promote cohesion establishment depends on multiple interactions between ESCO2 and the DNA replication machinery (Song et al., 2012; Bender et al., 2020; Ivanov et al., 2018; Moldovan et al., 2006). The relatively unstructured N terminal tail of ESCO2 contains several conserved motifs that promote interaction of ESCO2 with both the MCM2-7 replicative helicase and with proliferating cell nuclear antigen (PCNA), which forms a homotrimeric sliding clamp. Previously, interaction of ESCO2 with PCNA was shown to promote localization of ESCO2 at replication foci, cytologically detectable sites of active DNA replication, and this localization was shown to correlate with the ability to promote cohesion establishment (Bender et al., 2020). Precisely how interactions between ESCO2 and the MCM2-7 complex are managed in the context of active DNA replication, and how they are integrated with the formation and function of replication foci, is unclear. The mechanisms underlying the formation and function of replication foci are poorly understood; the cohesin complex contributes to the efficiency of DNA replication at these sites, perhaps through the formation of chromatin loops (Guillou et al., 2010).

Here we show that ESCO2 function is tightly regulated in a cell-cycle-dependent manner through a single post-translational modification in its N terminal tail. We show that ESCO2 modification impacts ESCO2’s ability to promote cohesion and alters interaction with the replication machinery. These data elucidate a mechanism that ensures that cohesion between sister chromatids is established in a cell cycle dependent manner, at least in part by affecting the subnuclear localization of a critical cohesion establishment factor.

## Results

### Impact of Specific Residues on Sister Chromatid Cohesion

The ESCO2 acetyltransferase modifies the SMC3 subunit of cohesin during S phase, and this activity is required for cohesion between sister chromatids. The interaction of ESCO2 with the MCM replicative helicase is essential for full ESCO2 activity (Bender et al., 2020; Ivanov et al., 2018; Higashi et al., 2012). This is consistent with the observation that ESCO2 recruitment to chromatin requires replication licensing, defined as the recruitment of the MCM helicase to chromatin (Higashi et al., 2012; Song et al., 2012). Although the MCM helicase is composed of six subunits, MCM2-7, the data suggest that ESCO2 interacts primarily with the MCM4 and MCM7 subunits (Ivanov et al., 2018). Interaction with other subunits of the complex remains possible. In ESCO2, the interaction has been mapped to a conserved motif called Box A in the N terminal tail of ESCO2 (Fig. 1A) (Ivanov et al., 2018; Song et al., 2012; Higashi et al., 2012; Bender et al., 2020).

**Figure 1.**
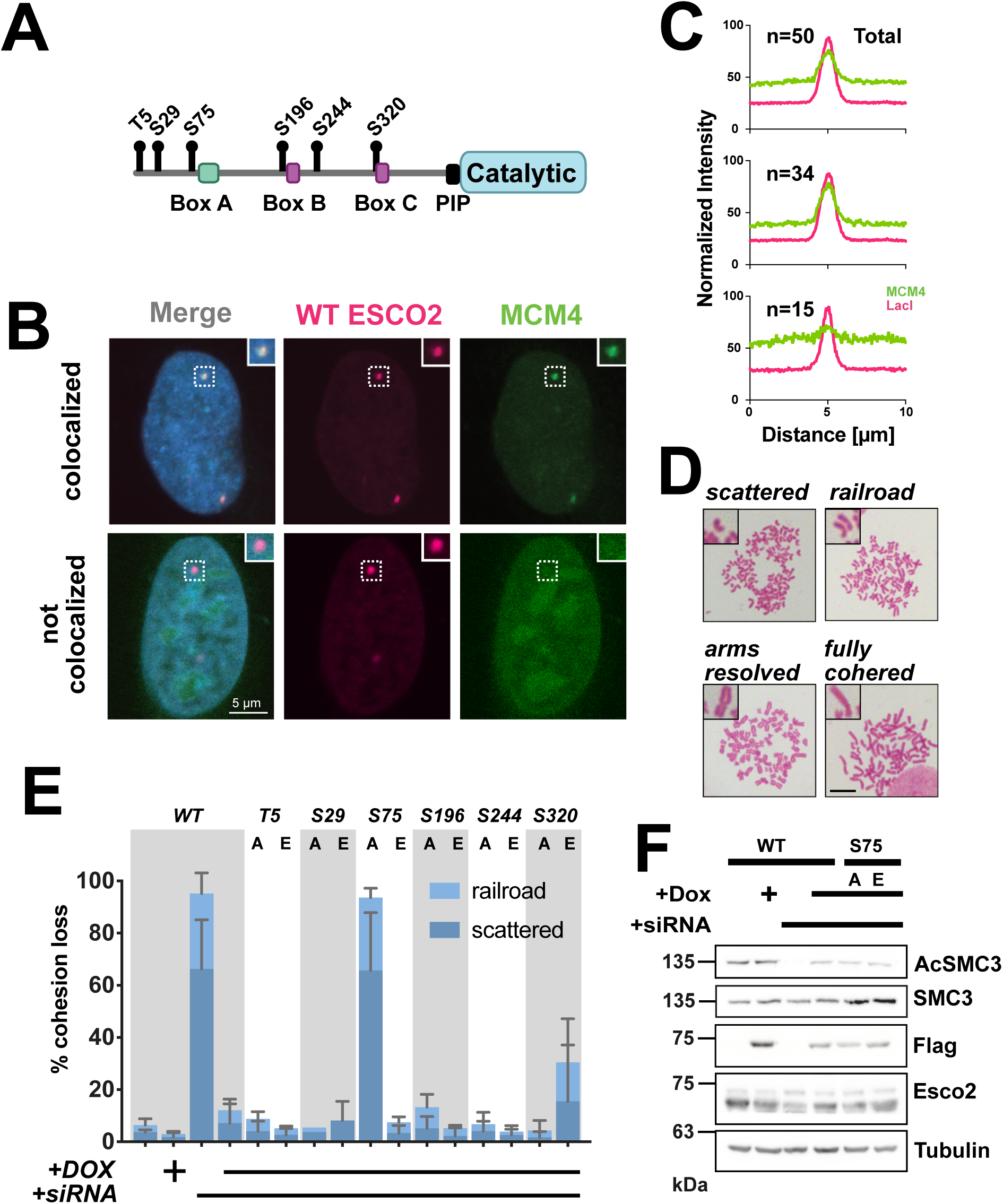
The impact of post-translational modification on ESCO2 function. **A. Graphic representation of ESCO2 protein.** Shown is a cartoon diagram of the ESCO2 protein. Several conserved motifs found in its N terminal unstructured tail have been identified and characterized. Small lollipops indicate potential phosphorylation sites in ESCO2, assayed below. **B. Colocalization of ESCO2 and MCM4 in cell populations.** Immobilized ESCO2 recruits MCM4 in most but not all cells. Confocal images of U2OS cells with a stably integrated array of *lac* operators transfected with constructs encoding mCherry-lacI-ESCO2 and GFP-MCM4. Images are representative of different phenotypes, in which MCM4 is or is not recruited to the lac array focus are shown (scale bar = 5 µm)**. C. Distinct phenotypes.** Normalized fluorescence intensity profiles of lines drawn across the lac array foci from cells expressing mCherry-lacI-ESCO2 and GFP-MCM4 could be manually sorted into high (middle) and low (bottom) colocalization phenotypes. Shown is a representative example of 3 independent experiments, n≥20. **D. Cohesion phenotypes.** Examples of phenotypes scored in analysis of mitotic chromosome spreads, in which scattered and railroad phenotypes are considered scored as cohesion loss. (scale bar = 10 µm) **E. Mutation of ESCO2 S75 alters ESCO2 function.** HeLa ESCO1^KO^ cells expressing siRNA-resistant derivatives of ESCO2 with indicated mutations were treated with siRNA against ESCO2 and chromosome spreads were scored for cohesion as illustrated in D. n ≥ 100 mitotic cells for all samples. Error bars represent one standard deviation from three independent experiments. **F. ESCO2-dependent cohesin acetylation does not correlate with function.** Whole cell lysates prepared from samples treated as in E were probed with the indicated antibodies. Acetylation of SMC3 (top panel) was similar in cells expressing the indicated ESCO2 derivatives: WT, S75A, and S75E.

To characterize further the interaction between ESCO2 and the MCM complex, we used an assay in which ESCO2 fused to the E. coli *lac* repressor protein is co-expressed with fluorescently tagged MCM4 protein, in a cell line that has an array of *lac* operators stably integrated into the genome. Through the LacI moiety, mCherry-tagged ESCO2 is immobilized at the lac array where it forms a detectable fluorescent focus. Here we assessed whether mEmerald-MCM4 was recruited to the ESCO2 focus. As previously, we found that fluorescent MCM4 was recruited to the ESCO2 focus (Fig. 1B) (Bender et al., 2020). To quantify this interaction, we averaged fluorescence intensity profiles of both ESCO2 and MCM4 across the ESCO2 foci (Fig. 1C). While characterizing the interaction, we found that the interaction is clear when measured by average intensity across the foci, but that approximately 25% of the cells showed no clear recruitment of MCM4 to the ESCO2 foci (Fig. 1C).

We wondered whether post-translational modification of ESCO2 might impact interaction with the replication machinery, and thus function. Post-translational modifications of ESCO2 have been identified in high-throughput screens, but data about the impact of these modifications on ESCO2 function was not available. Six potentially phosphorylated residues in ESCO2 were identified using the PhosphositePlus aggregator tool (Hornbeck et al., 2015), and these residues were mutated individually to either glutamic acid, to act as a phosphomimetic, or to alanine, to prevent phosphorylation (Fig. 1A). Using a knockdown and replacement strategy, the impact of the potential ESCO2 phospho-mutations on the ability of ESCO2 to promote cohesion establishment was assessed. Utilizing a cell line in which the related ESCO1 gene was knocked out, as previously (Alomer et al., 2017), endogenous ESCO2 was depleted with siRNA, and expression of siRNA-resistant ESCO2 derivatives was induced with doxycycline. All ESCO2 derivatives showed similar levels of expression (Supplemental Fig. 1). Mitotic chromosome spreads were prepared and sister chromatid cohesion was assessed (Fig. 1D, E). As previously, expression of siRNA-resistant wild type ESCO2 fully restored cohesion to control levels. Of the twelve mutations tested, only the ESCO2 derivative containing the S75 to alanine (S75A) mutation failed to promote cohesion establishment. Interestingly, the substitution of a phosphomimetic residue at the same site (S75E) restored normal levels of sister chromatid cohesion. We conclude from this experiment that phosphorylation of ESCO2 at S75 likely promotes the establishment sister chromatid cohesion, while the absence of phosphorylation at this residue limits cohesion establishment.

Interestingly, we saw little correlation between the level of SMC3 acetylation and the level of cohesion establishment in the assay shown in Figure 1E. The amount of SMC3 acetylation was similar in cells expressing wildtype, S75A, and S75E ESCO2 (Fig. 1F). We imagine that the context in which acetylation occurs may be critically important, and that cohesin acetylation can occur in non-productive ways. This effect has been seen previously, in related experiments (Song et al., 2012; Higashi et al., 2012)

### ESCO2 S75 modification impacts interaction of ESCO2 with MCM4

To investigate how phosphorylation of ESCO2 at S75 affects cohesion establishment, we tested whether the relevant mutations impact interaction between ESCO2 and the replication machinery. Because S75 is near the Box A MCM interaction motif (Fig. 1A), we first tested the impact of S75 modification on MCM interaction. Using the same recruitment assay shown in Figure 1A and B, we tested whether mutation of S75 impacted the ability of ESCO2 to interact with MCM4. In this assay we found that immobilized ESCO2 S75E was able to strongly recruit MCM4. In contrast, the non-phosphorylatable S75A mutant failed to recruit MCM4 (Fig.2A and B). These data are consistent with a model in which interaction between ESCO2 and the MCM complex is regulated by phosphorylation of ESCO2 at S75. To better quantify the recruitment, we measured assessed colocalization of ESCO2 with MCM4 at the foci by measuring the Pearson correlation coefficient between the two fluors in small regions of interest including the foci (Fig. S2). Consistent with the phenotype seen in Figure 1B, the Pearson correlation between immobilized ESCO2 and MCM4 appears bimodal, with some cells showing a low correlation and most cells showing a stronger correlation (Fig. 2C, green). In contrast, the correlation of ESCO2-S75E with MCM4 was higher than that of WT ESCO2, and the correlation between ESCO2-S75A and MCM4 was lower. We conclude that phosphorylation of ESCO2 at S75 promotes interaction of ESCO2 with MCM4, and assume that the bimodal colocalization seen with WT ESCO2 may reflect differences in its phosphorylation at S75 within a cell population.

**Figure 2.**
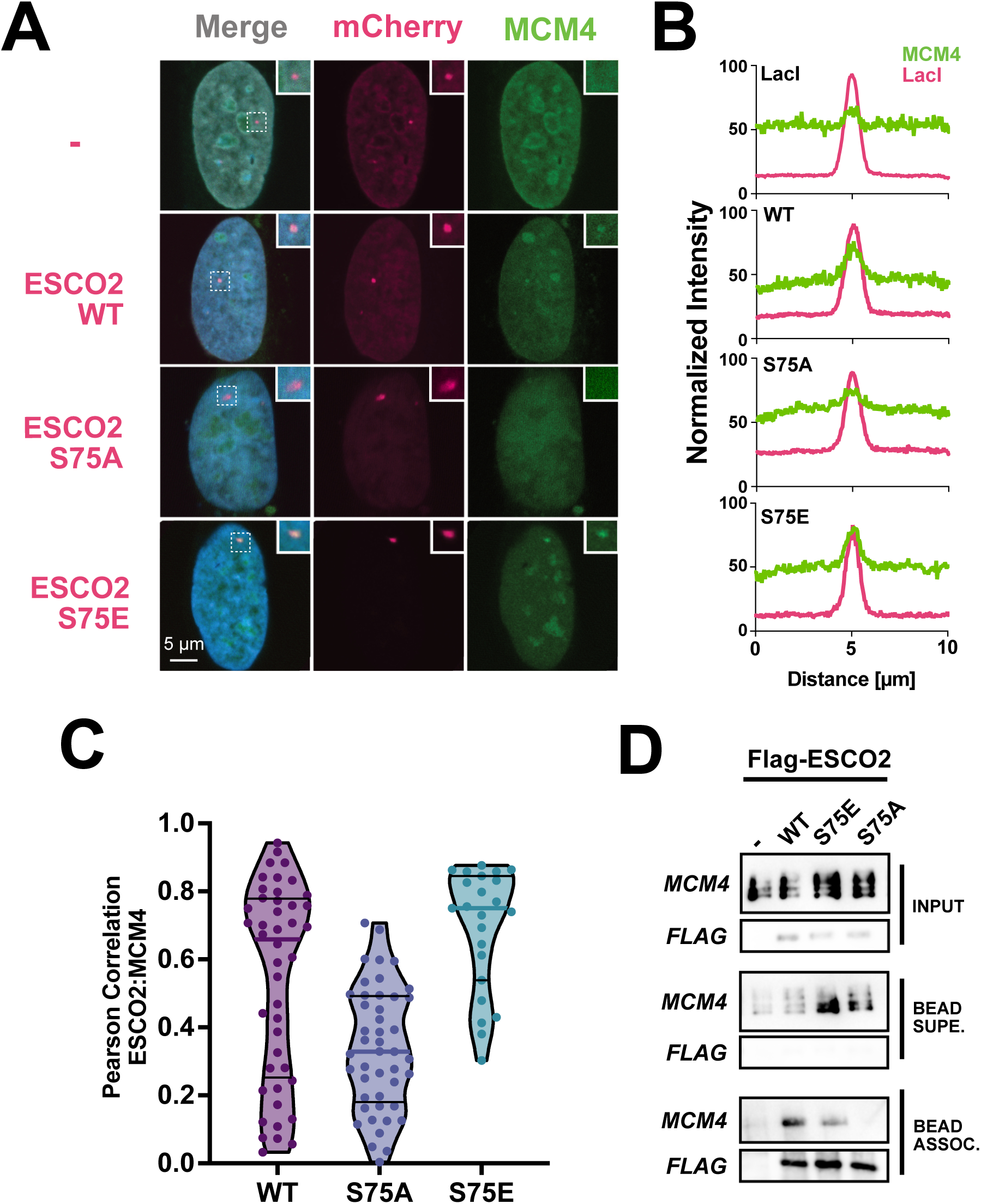
ESCO2 S75 impacts interaction of ESCO2 with MCM4. **A. Immobilized S75A ESCO2 has an impaired ability to recruit MCM4.** Confocal images of U2OS cells with immobilized lacI-fused S75 phosphomutant derivatives of ESCO2 were analyzed to assess the recruitment of mEmerald-MCM4, as in Figure 1. Tethered ESCO2 S75A showed a reduced ability to recruit MCM4 to the lac array compared to WT. Recruitment of MCM4 was unaffected in cells expressing ESCO2 S75E. (scale bar = 5 µm). **B. Normalized Fluorescence Intensity.** Profile lines of fluorescent intensity of mCherry-ESCO2 and GFP-MCM4 across the lac array were averaged and normalized, n≥ 15 cells. **C. ESCO2 S75A has reduced correlation with MCM4.** Violin plot of the Pearson correlation of mCherry-ESCO2 phosphomutant derivatives and GFP-MCM4. Each dot represents a measurement taken from a ROI around a lac array focus. **D. Coimmunoprecipitation of MCM4 with ESCO2.** Extracts of chromatin-associated proteins were prepared from HeLa cells expressing the indicated FLAG-tagged ESCO2 derivatives and synchronized in S phase. Following nuclease treatment, the solubilized samples were subjected to anti-FLAG immunoprecipitation and analyzed by immunoblot for the indicated proteins.

To confirm the dependency of the interaction with MCM4 on ESCO2 modification, we performed a co-precipitation experiment. HeLa cells expressing Flag-tagged wildtype, S75E, or S75A ESCO2 derivatives were synchronized in S phase. Chromatin-associated proteins were extracted, ESCO2 was immunoprecipitated directly from this mixture with anti-flag beads, and bead-associated proteins were analyzed for the presence of MCM4 protein. Endogenous MCM4 co-immunoprecipitated with both wildtype ESCO2 and ESCO2 S75E, but not with ESCO2 S75A (Fig. 2D). Interestingly, the S75E mutant appeared to precipitate MCM4 most efficiently, consistent with the model that phosphorylation of ESCO2 at S75 promotes interaction with the MCM complex, and suggesting that the wildtype protein may have been only be partially phosphorylated in the cell lysate.

### Phosphomimetic ESCO2 has impaired localization to replication foci

We previously showed that ESCO2 colocalizes with PCNA at sites of active DNA replication called replication foci. Localization of ESCO2 to these foci depends on interactions between ESCO2 and PCNA, which occur through several PCNA-interacting protein boxes (PIP boxes) in the tail of ESCO2 (Bender et al., 2020). One PIP box in particular (a.a. 325-330 in the human protein), which we called Box C, is essential for both ESCO2 localization at replication foci and for cohesion establishment, suggesting that localization at RF is essential for full function (Bender et al., 2020).

How ESCO2 interactions with the MCM helicase and PCNA are coordinated, and whether they occur sequentially or simultaneously remains unresolved. To clarify this issue, we next determined whether ESCO2 phosphorylation impacts localization of ESCO2 to replication foci. To do this, mCherry-tagged ESCO2 derivatives were co-expressed in U2OS cells with GFP-tagged PCNA. As previously, wild type ESCO2 colocalized with PCNA in replication foci (Fig. 3A). Similarly the non-phosphorylatable S75A derivative also localized to replication foci.

**Figure 3.**
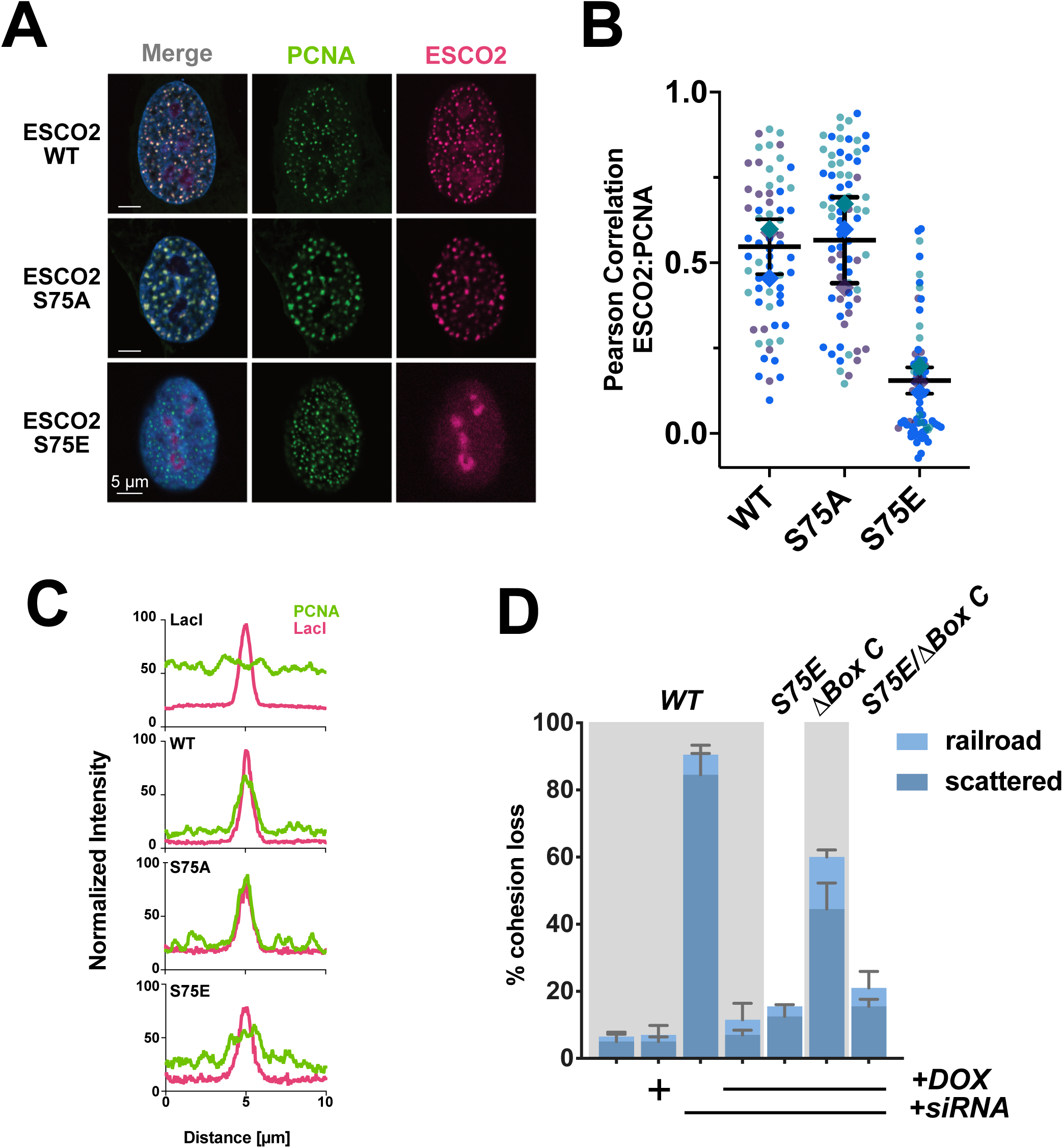
Altered interaction of mutant ESCO2 with replication foci **A. ESCO2 S75E show reduced localization at replication foci**. Confocal images of U2OS cells expressing mCherry-lacI-ESCO2 fusion, with indicated mutations in ESCO2, and a GFP-PCNA fusion. Sites of colocalization appear yellow in merged images. ESCO2 S75E has impaired localization with replication foci and accumulates in the nucleolus. (scale bar = 5 µm). **B. ESCO2 S75E has reduced correlation with nuclear PCNA.** Pearson correlation between ESCO2 and PCNA was calculated from images as shown in A. Each dot represents a measurement taken from a single ROI within a nucleus, excluding nucleoli. Shown is a Superplot of measurements from three independent experiments indicated by color, n≥20 cells. Error bars represent the standard deviation of the averages of the three experiments. **C. Immobilized ESCO2 S75E has an impaired ability recruit PCNA to nuclear foci.** Fluorescent foci of immobilized mCherry-lacI-ESCO2 in U2OS cells (as in Fig. 1 and 2) were analyzed for colocalization of PCNA, as previously. **D. Mutation at S75 suppresses phenotype of Box C deletion.** Shown is a cohesion assay, as in Figure 1, in which the endogenous ESCO2 is depleted and ESCO2 transgenes are induced with doxycycline. While the Box C deletion only partially rescues cohesion, the double mutant, S75E/ΔBox C more fully restores cohesion.

Strikingly, however, the phosphomimetic mutant, S75E, failed to colocalize with PCNA in replication foci (Fig. 3A, bottom). As seen previously, the ESCO2 protein that was not associated with replication foci accumulated instead in the nucleoli (Bender et al., 2020). To quantify colocalization between ESCO2 and PCNA, we measured the Pearson correlation between the fluorescence intensities of ESCO2 and PCNA in the nuclei (Fig. 3B). To do this, the correlation between ESCO2 and PCNA in non-nucleolar regions (selected in the DAPI channel) was measured within a subnuclear region. Consistent with the visual assessment, the S75E mutant showed a much lower colocalization with PCNA than either the wild type or the S75A mutant. We draw two conclusions from this experiment: first that phosphorylation of ESCO2 at S75 blocks enrichment at replication foci, and secondly, that overt enrichment at replication foci is not strictly required for cohesion establishment by ESCO2.

At this point we imagined two possible explanations for the impact of ESCO2 phosphorylation on cohesion establishment. In one model, interaction of ESCO2 with PCNA and the MCM helicase would be mutually exclusive, and ESCO2 phosphorylation would result in constitutive interaction with the MCM helicase. In a second model, S75 phosphorylated ESCO2 might still interact with PCNA, but fail to localize to replication foci for other reasons. To determine which of these two models is likely to be true, we again utilized the fluorescent two-hybrid assay shown in Figures 1 and 2, immobilizing ESCO2 at the lac array, and expressing GFP-PCNA in the same cells (Fig. 3C). As previously, PCNA was strongly recruited to interact with both wildtype and S75A ESCO2. In contrast, recruitment of PCNA by ESCO2 S75E was reduced suggesting that phosphorylation at S75 may weaken the interaction between ESCO2 and PCNA. This may suggest that forced interaction with the MCM helicase alone is sufficient to ensure cohesion establishment, and that localization to replication foci normally ensures that ESCO2 is somehow presented appropriately to the MCM complex.

What is the evidence that association with replication foci is important for ESCO2 function? We previously showed that the PIP box in ESCO2 called “Box C” is required for both cohesion establishment and localization of ESCO2 to replication foci. However our new data showing that the S75E mutant reduces interaction of ESCO2 with PCNA but fully rescues cohesion led us to test whether the ΔBox C defect in cohesion establishment could be suppressed by the S75E mutation. If so, it would suggest a model in which forced association with MCM obviates the need for high affinity interaction with PCNA. We therefore generated a double mutant containing both the Box C deletion with the phosphomimetic S75E mutation and assessed its ability to promote cohesion using the gene knockdown and replacement strategy as in Figure 1.

Consistent with previous results, the ESCO2 ΔBox C mutant showed a reduced ability to promote sister chromatid cohesion, and the ESCO2 S75E mutant fully restored cohesion to wild type levels (Fig. 3D). Finally, the double mutant, S75E/ΔBox C fully rescued cohesion, suggesting that stable interaction with the MCM helicase is sufficient for ESCO2’s role in cohesion establishment, even when the strongest PCNA interacting motif is deleted. It remains possible that lower affinity interactions with PCNA through Box B and the canonical PIP box may play roles without being sufficient to alter subnuclear localization.

### ESCO2 is a CDK2 substrate

The consensus sequence flanking ESCO2 S75 suggests that it might be a target of cyclin-dependent kinases (CDKs) (Fig. 4A). CDKs are a well characterized family of serine/threonine kinases that are activated by association with cyclin subunits and preferentially phosphorylate substrates at [S/T]P-K/R motifs (Stevenson-Lindert et al., 2003; Suzuki et al., 2015; Songyang et al., 1994). Because multiple CDKs regulate cell cycle progression, we sought to determine which cyclin/CDK complex most likely phosphorylates ESCO2. To this end, we again immobilized ESCO2 at the *lac* array and analyzed the impact of selective CDK inhibitors on recruitment of MCM4. With the addition of Ro-3306, a CDK1 inhibitor, recruitment of MCM4 to the *lac* array was unaffected (Fig. 4B). In contrast, when the assay was performed with NU6102, a CDK1/2 inhibitor, MCM4 recruitment was reduced. This suggests that CDK2 activity promotes interaction between ESCO2 and the MCM helicase. We were unable to identify an effective CDK2-specific inhibitor.

**Figure 4.**
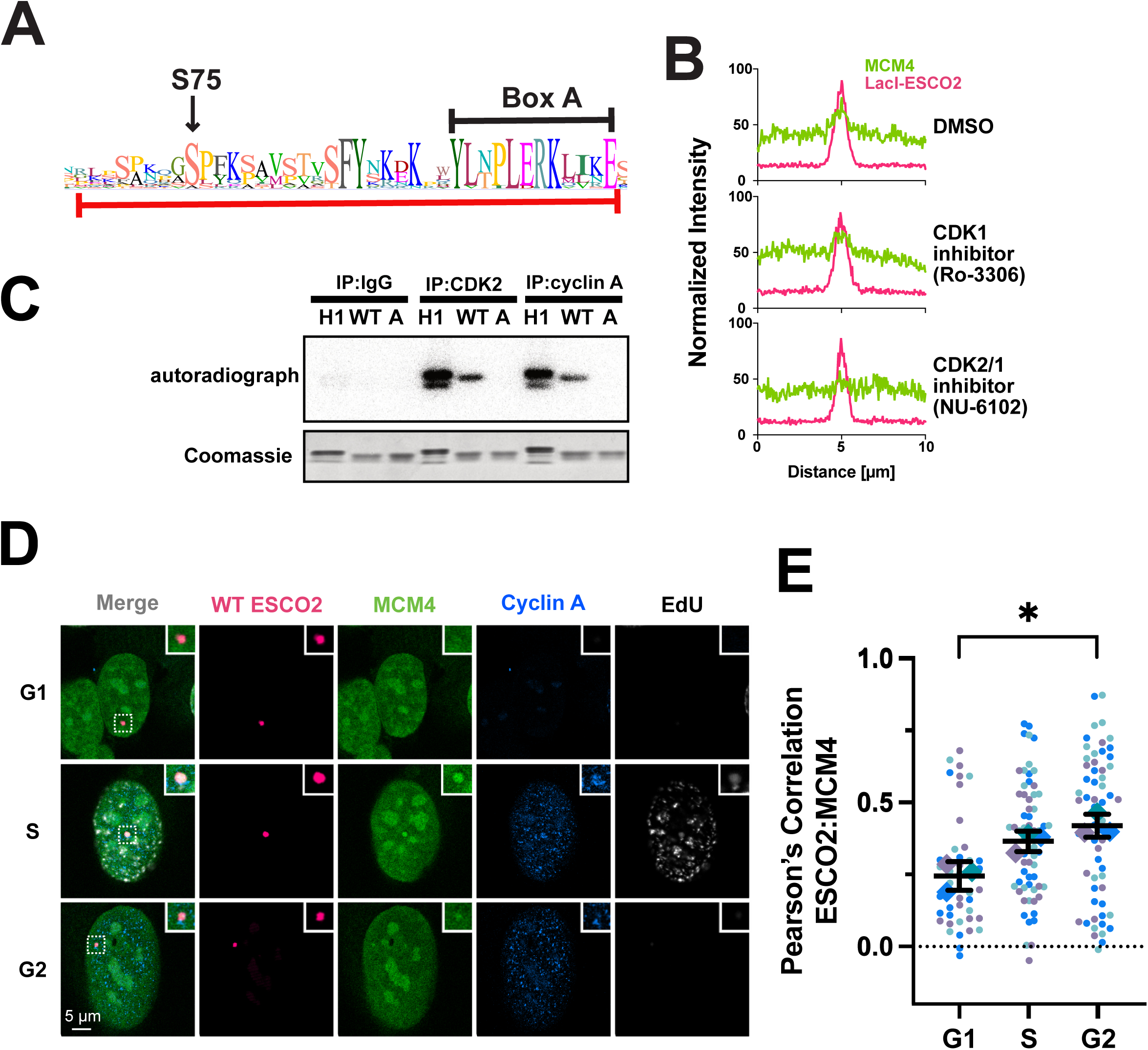
ESCO2 is a CDK2 substrate **A. Sequence conservation around ESCO2 S75.** Shown is a sequence logo, in which the degree of conservation at each position is indicated by the size and height of the letter that indicates the amino acid. ESCO2 S75 is indicated, as well as the details of Box A illustrated in Figure 1. The red line indicates the sequence used in the assay shown in C. **B. The impact of Cdk inhibition on the ESCO2-MCM4 interaction.** Immobilized ESCO2 foci were scored for recruitment of MCM4 as in Figure 1 and 2, this time in the presence of the indicated inhibitors. DMSO was used as a vehicle control, n= 15 cells. **C. ESCO2 is phosphorylated at S75 by Cdk2.** Immunoprecipitated CDK2 or Cyclin A were incubated with linker histones (H1) or recombinant ESCO2 fragments fused to GST in the presence of [γ-32P] ATP. The fused ESCO2 peptide sequence is indicated by the red line in **A**. Both wild type and S75A ESCO2 derivatives were tested, as indicated. The reaction mix was resolved by SDS-PAGE and labeled proteins were detected by phosphor-imager autoradiogram (top) and Coomassie staining (bottom). **D. ESCO2 and MCM4 colocalize in correlation with Cyclin A expression.** Confocal images of U2OS cells with a stably integrated array of *lac* operators transfected with constructs encoding mCherry-lacI-ESCO2 and GFP-MCM4. Images are representative of different stages of the cell cycle, identified by EdU and cyclin A expression. (scale bar = 5 µm) **E. ESCO2 correlation with MCM4 increases as cell cycle progresses.** Pearson correlation between ESCO2 and MCM4 was calculated from images as shown in D. Each dot represents a measurement taken from a single ROI drawn around a lac array. Shown is a Superplot of measurements from three independent experiments indicated by color, n≥50 cells.

We next used a purified system to confirm that ESCO2 can serve as a substrate of CDK2-dependent phosphorylation. To do this we purified a recombinant ESCO2-derived 38a.a. peptide fused to glutathione S transferase (GST) to use as a substrate. This peptide included the S75 residue as well as adjacent residues (indicated by the red line in Fig. 4A). We then immunoprecipitated CDK2 from a lysate of asynchronously growing cultured cells, and incubated the precipitated kinase with the recombinant GST fusion protein in the presence of radiolabeled [γ-32P]ATP. As a positive control, we used histone H1 as a substrate. Reactions were resolved by SDS-PAGE, and phosphorylation was detected by autoradiography (Fig. 4C). A radioactive signal consistent with phosphorylation of the ESCO2 fusion was observed. Importantly, this reaction product was not detected when the recombinant substrate contained the S75A substitution, confirming the specificity of phosphorylation of ESCO2 at S75 by CDK2.

Similarly, a Cyclin A immunoprecipitate was able to phosphorylate the GST-ESCO2 fusion in an S75-dependent manner. Together these data are consistent with a model in which ESCO2 is phosphorylated by CyclinA/CDK2 at S75. It remains possible that Cyclin E/CDK2 might also modify ESCO2 at S75; we were unable to obtain an effective anti-Cyclin E antibody to test this model directly.

Our data suggest that ESCO2 activity is critically dependent on CyclinA/Cdk2, which promotes the interaction of ESCO2 with the MCM complex. This model predicts that ESCO2 should interact with MCM proteins only during or after DNA replication, as cyclin A levels are very low in G1, and rise during S phase, peaking in G2 (Elledge et al., 1992; Arooz et al., 2000). To test this, model, we again analyzed interaction of ESCO2 with MCM4 using the fluorescent 2-hybrid assay with cells expressing tethered ESCO2 (wild type) and GFP-tagged MCM4, in this case combining the assay with immunostaining for cyclin A and pulse labeling of the cells with the thymidine analog EdU to identify cells undergoing DNA replication. With these data were able to assign interphase cells into three categories: G1 (cyclin A negative, EdU positive) S phase (Cyclin A positive, EdU positive) and G2 (cyclin A positive, EdU negative) (Fig. 4C). Within each population we measured the correlation between signals from immobilized ESCO2 and coexpressed MCM4. Consistent with our model, interaction of MCM4 with ESCO2 at the lacO array was lowest in G1, increased in S phase, and was highest in G2(Fig. E).

## Discussion

Here we have shown that ESCO2, which is essential for cohesion between sister chromatids, is a direct target of cell cycle control. CDK-dependent modification is critical to the ability of ESCO2 to promote cohesion during DNA replication. It also modulates association of ESCO2 with the replication machinery (Fig. 5).

**Figure 5.**
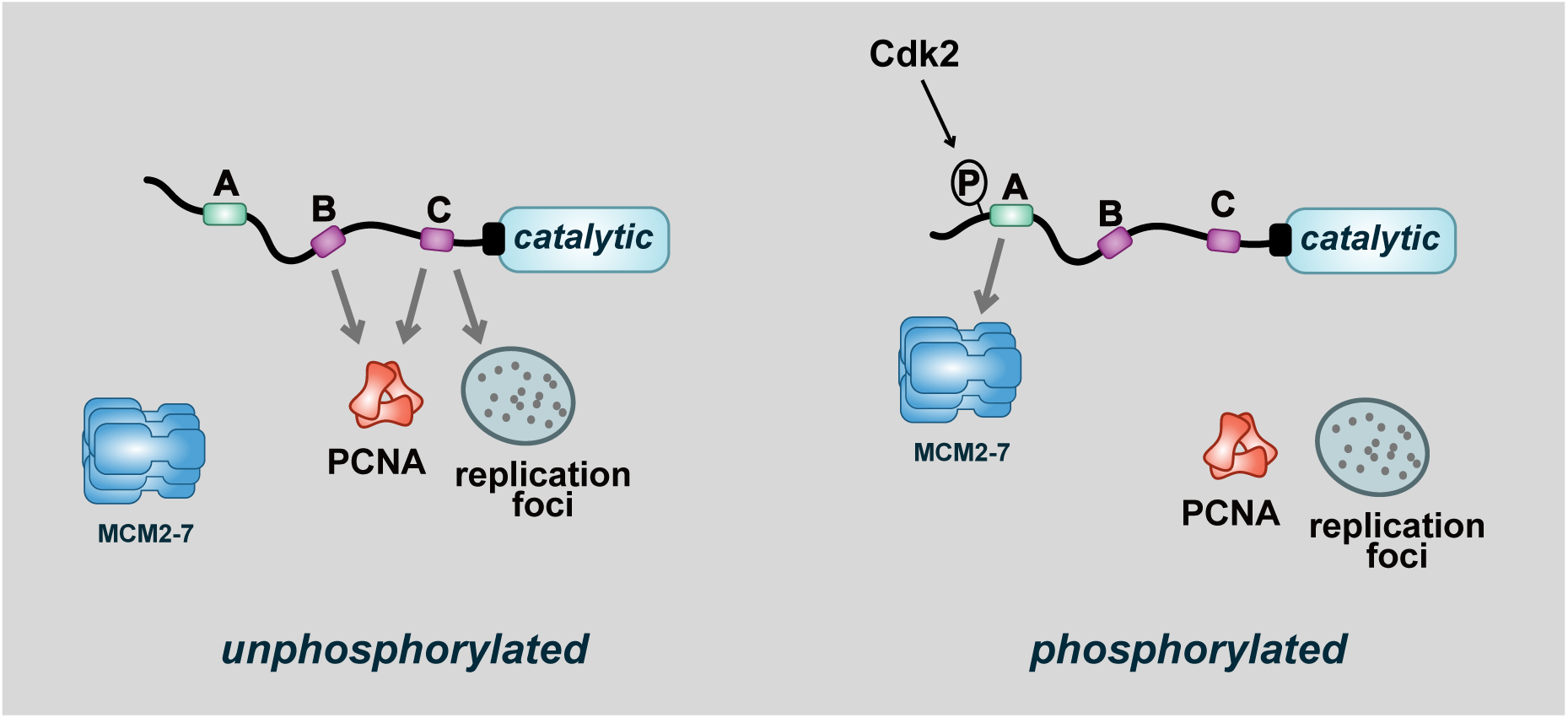
Model: CDK2-dependent phosphorylation alters ESCO2 association with the replication machinery. **Left**: in the absence CDK2 activity, unphosphorylated ESCO2 interacts with PCNA and is concentrated in replication foci. **Right**: CDK2-dependent phosphorylation at S75 increases association of ESCO2 with the MCM complex and promotes cohesion establishment.

The activity of the cohesin complex is tightly controlled during cell cycle progression. Prior work has shown that the association and engagement of cohesin with chromatin is controlled at several levels by cell cycle-dependent mechanisms. The interaction of the cohesin loader, SCC2/SCC4 with chromatin is integrated with replication licensing and replication initiation, which is cell cycle controlled (Takahashi et al., 2008, 2004; Gillespie and Hirano, 2004). The removal of cohesin from chromatin in prophase, particularly in vertebrates, is dependent upon cell cycle-dependent phosphorylation of cohesin subunits, and chromosome separation at anaphase depends both on phosphorylation-dependent removal of cohesin and Sororin from chromatin, and proteolytic cleavage of Securin (Zou et al., 1999; Hauf et al., 2001; Waizenegger et al., 2000; Cohen-Fix et al., 1996; Dreier et al., 2011; Uhlmann et al., 1999). Precisely how cohesion establishment is coupled with DNA replication has been elusive. Our work here adds a new detail to this literature. Here we have shown that CDK2 phosphorylates ESCO2, and this phosphorylation is required for ESCO2 to promote cohesion establishment.

ESCO2 interacts with proteins associated with active replication, including PCNA and the MCM helicase. How these interactions are coordinated and integrated with DNA replication, and whether they occur simultaneously, has been unclear (Bender et al., 2020; Higashi et al., 2012; Ivanov et al., 2018; Moldovan et al., 2006). During cell cycle progression the loading of the MCM complex and PCNA are temporally distinct. The MCM complex is loaded, in excess, in G1 in a step called replication licensing, whereas PCNA is loaded later, behind active replication forks on both the leading and lagging DNA strands (Evrin et al., 2009; Remus et al., 2009; Hyrien, 2016; Kirstein et al., 2021). Because ESCO2 is a substrate of APC-dependent degradation in G1, its levels are very low in G1 (Song et al., 2012; Lafont et al., 2010). It accumulates in S phase, when it associates with chromatin, and can then interact with the MCM complex and/or PCNA. The data presented here suggest that interaction with the MCM complex and PCNA may be mutually exclusive: phosphorylated ESCO2 interacts strongly with the MCM complex, while unmodified ESCO2 preferentially interacts with PCNA, and no longer effectively binds the MCM complex. Both CDK and phosphatase activities have been known to drive critical steps in DNA replication, including firing of replication origins at initiation and removal of initiation factors (Jenkinson et al., 2023; Masumoto et al., 2002; Tanaka et al., 2007; Zegerman and Diffley, 2007; Sansam et al., 2015; Kumagai et al., 2011; Boos et al., 2011). Here we have shown that CDK activity is important for replication-associated cohesion establishment. It will be interesting to determine whether dephosphorylation of ESCO2 S75 plays a role in cohesion establishment.

But how does CDK-dependent phosphorylation happen in the context of DNA replication? Where does phosphorylation of ESCO2 occur relative to fork progression? In previous work, we found a correlation between the ability of ESCO2 to promote cohesion and its enrichment at the replication foci (Bender et al., 2020). Here, however, we have found that ESCO2 phosphorylation likely prevents the subnuclear localization of ESCO2 to replication foci, even though the phosphomimetic derivative is fully able to promote cohesion establishment. These observations suggest a model in which replication foci provide an environment that promotes or enhances ESCO2 phosphorylation, which in turn promotes ESCO2 association with the MCM complex. In this model, the S75E mutation, by mimicking phosphorylation, would bypass the requirement for association with replication foci. Importantly, the combination of the phosphomimetic S75E mutation with the box C deletion fully restores cohesion establishment, consistent with the model that association of ESCO2 with replication foci in some way promotes its phosphorylation. Stable interaction of ESCO2 with the MCM complex is sufficient for establishment of sister chromatid cohesion. It is also possible that interactions with replication foci and the MCM complex provide redundant pathways for cohesion establishment.

There is an emerging consensus that cohesion between sister chromatids can be established by one of two different pathways (Oldenkamp and Rowland, 2024). In the first, DNA is entrapped by cohesin prior to DNA replication, and cohesin is “converted” to holding two sisters together by the event(s) of DNA replication, or replication termination (Cameron et al., 2024; Srinivasan et al., 2020). In the second, the “de novo” pathway, cohesion is established behind the replication fork, likely through a single-stranded DNA intermediate, causing the two sister chromatids to be entrapped by the cohesin ring as they are formed (Murayama et al., 2024). It is interesting to speculate about the pathway to which ESCO2 contributes. ESCO2 may remain associated with the MCM helicase as the fork progresses, acetylating and stabilizing cohesin before the replisome passes (conversion pathway), or it may interact with PCNA behind the replication fork, contributing to the *de novo* pathway. Association of ESCO2 with the replication machinery suggests that ESCO2 is most likely to be involved in conversion, consistent with previous data suggesting synthetic lethality between loss of ESCO2 and of the helicase DDX11 (Faramarz et al., 2020; Abe et al., 2016). Since our data suggest that interaction of ESCO2 with replication foci is dispensable for cohesion establishment once ESCO2 is phosphorylated, we posit that accumulation at replication foci most likely serves to increase the effective local concentration of ESCO2 near sites of active replication, but is not critical for function. Association of ESCO2 with nascent DNA has been documented previously, consistent with continued association with the MCM complex during DNA replication (Alabert et al., 2014).

## Materials and Methods

### Cell Culture and Cell Line Construction

Cells were routinely cultured in Dulbecco’s modified Eagle’s medium (DMEM, Corning) supplemented with 10% fetal bovine serum (FBS, R&D Systems) and maintained at 37°C in 5% CO_2_ atmosphere. Stable cell lines with a doxycycline-inducible siRNA-resistant flag-tagged ESCO2 transgene were generated using HeLa Flp-In T-Rex ESCO1 KO cells as previously (Alomer et al., 2017). To do this, an siRNA resistant derivative of an ESCO2 cDNA (the sequence CGAGTGATCTATAAGCCAA was modified to CGTGTCATTTACAAACCTA) was cloned into a pcDNA5/FRT-based flag-tagged vector and that plasmid was cotransfected with a plasmid expressing the FLP recombinase (pOG44, Invitrogen) using Lipofectamine 3000 (Invitrogen) according to the manufacturer’s instructions. Cells were incubated overnight and then sparsely plated in media with 200 μg/mL hygromycin B (Gold Biotechnology) for 15 days. Individual colonies were collected with trypsin-EDTA (Invitrogen) soaked filter paper and amplified. Transgene expression was induced by overnight incubation in 2 μg/mL of doxycycline (VWR) and confirmed by immunoblot.

### Cohesion Assays

HeLa Flp-In T-Rex ESCO1^KO^ cells with doxycycline-inducible siRNA-resistant flag-tagged ESCO2 transgene stably integrated were treated with 2 μg/mL of doxycycline(VWR) for 24 hours to induce expression of the ESCO2 derivative. Endogenous ESCO2 was depleted by transfecting cells with 20nM siRNA (Dharmacon, J-025788-09, target: CGAGUGAUCUAUAAGCCAA) using Lipofectamine RNAiMAX (Invitrogen) according to the manufacturer’s instructions. The cells were incubated at 37° C in 2 µg/mL doxycycline for 42 hours. Cells were collected in trypsin-EDTA (Invitrogen), washed with PBS, and treated with a hypotonic solution (75 mM KCl) for 20 minutes at room temperature. Cells were pelleted, resuspended in 200 µL of hypotonic solution, and fixed in 5 mL of ice cold 3:1 methanol:acetic acid. Samples were resuspended twice in fresh fix and dropped onto glass slides. The slides were stained with Giemsa (VWR) and coverslips were mounted with Permount (Fisher). Images were collected using a Zeiss Axio Imager Z. 1, using a 63x 1.4 n.a. oil immersion lens. Phenotypes were scored blind in three biological replicates and at least 100 cells were scored per sample.

### Co-immunoprecipitation Assays

HeLa Flp-In T-Rex ESCO1^KO^ cells with doxycycline-inducible siRNA-resistant flag-tagged ESCO2 transgene stably integrated were treated with 2 μg/mL of doxycycline for 24 hours to induce expression of the ESCO2 derivative. Cells were arrested for 20 hours with 2 mM thymidine (Sigma Aldrich) and released into S phase for 2.5 hours. Cells were trypsinized, washed in PBS, and resuspended lysis buffer (75 mM HEPES pH 7.5, 1.5 mM MgCl_2_, 150 mM KCl, 15% glycerol, 1.5 mM EGTA pH 8.0, 0.075% NP40, 1 mM DTT, 0.1 mM PMSF, 10 µg/mL leupeptin, 10 µg/mL chymostatin, 10 µg/mL pepstatin, 20 mM β-glycerophosphate, 10 mM NaF, 1 mM Na3VO4). Lysates were subjected to two freeze-thaw cycles, followed by douncing 20 times on ice. Soluble and chromatin-bound fractions were separated by centrifugation.

Chromatin pellets were washed five times with Wash Buffer I (50 mM HEPES pH 7.5, 1.5 mM MgCl2, 150 mM KCl, 10% glycerol, 1.5 mM EGTA pH 8.0, 0.075% NP40, 1 mM DTT, 0.1 mM PMSF, 10 µg/mL leupeptin, 10 µg/mL chymostatin, 10 µg/mL pepstatin, 20 mM β-glycerophosphate, 10 mM NaF, 1 mM Na3VO4), resuspended in Wash Buffer I containing 300 U/mL Benzonase, and rotated at 4°C for 2 hours. The reaction was centrifuged at 2,500 g for 2 min to and the supernatant containing chromatin-bound proteins was used for IP. Samples were incubated with anti-flag affinity resin (VWR) on a roller overnight at 4°C. Beads were washed four times with Wash Buffer I and proteins were eluted in sample buffer for analysis by western blot.

### Gels and Immunoblots

Protein samples were resolved on 7 to 15% gradient SDS-PAGE gels and transferred to nitrocellulose membrane using the Trans-Blot Turbo (Bio-Rad). Membranes were incubated in 5% milk in TBST for 1 hour at room temperature, probed with primary antibodies overnight at 4° C, washed three times with TBST, probed with horseradish peroxidase (HRP)-conjugated secondary antibodies for 45 minutes at room temperature, washed three times with TBST, and two times with TBS. Signals were detected with chemiluminecent substrate (Licor Biosciences) and imaged with the Azure 300 imager (Azure Biosystems).

### Immunofluorescence

For visualization of replication foci, U2OS cells were cotransfected with 500 ng of pCS2-GFP-PCNA and 500ng of mcherry-ESCO2 derivatives using Lipofectamine 3000 according to the manufacturer’s protocol. Cells were incubated on glass coverslips for 48 hours at 37°C in 5% CO_2_ atmosphere. The cells were washed in phosphate buffered saline (PBS), pre-permeabilized in ice-cold CSK buffer (10mM HEPES pH 7.4, 100 mM NaCl, 300 mM sucrose, 3 mM MgCl2, 0.05% Triton) on ice for 5 minutes to extract soluble proteins. The cells were washed again in 1X PBS and fixed in 2% PFA for 18 minutes. Cells were counterstained with DAPI and the coverslips were mounted onto slides. Images were collected on a Nikon C2 confocal on a Ti-E motorized inverted microscope using a 60x oil immersion objective. To analyze colocalization, NIS Elements software was used to draw a region of interest (ROI) within the nucleus excluding the nucleolus, and the Pearson correlation coefficient of the relevant fluorescence signals within the ROI was collected.

For immobilization assays, U2OS LacO-I-SceI-TetO cells (Kerafast) were cotransfected with 500ng of plasmids encoding mCherry-ESCO2 derivatives 250 ng of pCS2-GFP-PCNA or 250ng of pmEmerald-MCM4-C-19 using Lipofectamine 3000 according to the manufacturer’s protocol. Cells were incubated on glass coverslips for 48 hours at 37° C in 5% CO_2_ atmosphere. The cells were washed in PBS and fixed in PBS with 2% PFA and 0.1% Triton X-100 for 20 minutes at RT. Coverslips were washed with Abdil, stained with DAPI, and mounted onto slides. Images were collected as above. NIS Elements software was used to measure the fluorescence intensity along a 10 µm line. Pearson correlation between fluorescent proteins and immobilized ESCO2 was determined for regions of interest including and surrounding the tethered ESCO2 focus, using NIS Elements.

### Cell Cycle Image Analysis

U2OS LacO-I-SceI-TetO cells were transfected as previously. Cells were pulsed for 15 minutes with 20mM EdU (ThermoFisher), washed with PBS, and fixed in PBS with 2% PFA and 0.1% Triton X-100 for 20 minutes at RT. Coverslips were incubated for 30 minutes at RT in Click-iT reaction mix (CuSO_4_, Ascorbic Acid, Alexa Fluor™ 647 Click-IT azide (ThermoFisher), PBS).

Slips were washed in PBS and incubated overnight in cyclin A primary antibody (BioLegend) followed by 45 minutes of anti-mouse IgG 405 (Dylight). Images were collected on a Nikon C2 confocal on a Ti-E motorized inverted microscope using a 60x oil immersion objective.

Background-subtracted images were used to assess cyclin A staining, and cells were considered Cyclin A–positive if more than 1% of the total nuclear area exceeded a threshold assigned for each experiment. Thresholding to avoid nucleolar signal and include nuclear foci (around 250-350 intensity units in a range of 4095) was set once and used for all samples in a given experiment (Supp. Fig. 2). Cells that were negative for both Cyclin A and EdU were assigned to the G1 phase. EdU-positive cells were classified as S phase, while cells positive for Cyclin A but negative for EdU were designated as G2 phase. Pearson correlation between fluorescent proteins and immobilized ESCO2 was determined for regions of interest including and surrounding the tethered ESCO2 focus.

### IP Kinase Assay

Cell lysates were prepared using a HeLa cell line (ATTC). Cells were harvested with trypsin, washed with PBS, and reuspended in lysis buffer (75 mM HEPES pH 7.5, 1.5 mM MgCl2, 150 mM KCl, 15% glycerol, 1.5 mM EGTA pH 8.0, 0.075% NP40, 1 mM DTT, 0.1 mM PMSF, 10 µg/mL leupeptin, 10 µg/mL chymostatin, 10 µg/mL pepstatin, 20 mM β-glycerophosphate, 10 mM NaF, 1 mM Na3VO4). Cells were subjected to two freeze thaw cycles, dounced on ice, the supernatant was collected following centrifugation. For immunoprecipitation, 2 mg/mL antibody was added to 200 µL of cell lysates and incubated at 4°C for overnight. Protein A sepharose beads were washed 2-3 times with lysis buffer, and 40 µL of a 50% slurry of beads in lysis buffer was added to the lysates. After a 1-hour incubation at 4°C with rocking, beads were pelleted, and supernatant was removed. Beads were washed three times with 1 mL of bead wash buffer (1 M NaCl, 20 mM Tris pH 7.4) and once with 1x kinase reaction buffer (10 mM HEPES pH 7.5, 10 mM MgCl2).

The kinase reaction mixture contained 1x kinase reaction buffer, 6 µCi [γ-32P] ATP, 1 µL of 5 mM ATP and 5 mg/mL of substrate: Histone H1, GST-ESCO2 WT, or GST-ESCO2 S75A. Reactions were initiated by adding 5 µL of kinase cocktail to the immunoprecipitation beads and incubated at room temperature for 30 minutes. Reactions were terminated by adding an equal volume of 2x sample buffer with DTT, boiled for 5 minutes, and resolved by gel electrophoresis. Gels were dried, exposed to a phosphoimager screen for 24 hours and imaged on a Personal Molecular Imager (Bio-Rad).

**Table 1:** Plasmids used in this study.

| Plasmid Number | Plasmid Name | Source |
| --- | --- | --- |
| pAP11 | pcDNA5/FRT-TO-sirESCO2* | (Bender et al., 2020) |
| pAP14 | pcDNA5/FRT-TO-sirESCO2ΔC | (Bender et al., 2020) |
| pAP18 | pCS2/mCherry-LacI-NLS | (Bender et al., 2020) |
| pBR36 | pGEXCS/GST-ESCO2:67-104 | This work |
| pBR37 | pGEXCS/GST-ESCO2:67-104(S75A) | This work |
| pBR39 | pcDNA5/FRT-TO-3XFLAG-sirESCO2 S75E/ΔC | This work |
| pDB20 | pCS2/mcherry-LacI-NLS-ESCO2 | (Bender et al., 2020) |
| pDF01 | pCS2/mcherry-LacI-NLS-ESCO2/S75A | This work |
| pDF02 | pCS2/mcherry-LacI-NLS-ESCO2/S75E | This work |
| pJC260 | pcDNA5/3XFLAG-ESCO2 | This work |
| pOG44 | FLP recombinase | Invitrogen |
| pSR312 | pCS2/GFP-PCNA | Gift from Sansam Lab |
| (pSR349) | mEmerald-MCM4-C-19 | Addgene #54167 |
| pVM01 | pcDNA5/3FLAG-sirESCO2 S29A | This work |
| pVM02 | pcDNA5/3FLAG-sirESCO2 S29E | This work |
| pVM03 | pcDNA5/3FLAG-sirESCO2 S75A | This work |
| pVM04 | pcDNA5/3FLAG-sirESCO2 S75E | This work |
| pVM05 | pcDNA5/3FLAG-sirESCO2 S196A | This work |
| pVM06 | pcDNA5/3FLAG-sirESCO2 S196E | This work |
| pVM07 | pcDNA5/3FLAG-sirESCO2 S320A | This work |
| pVM08 | pcDNA5/3FLAG-sirESCO2 S320E | This work |
| pVM09 | pcDNA5/3FLAG-sirESCO2 T5A | This work |
| pVM10 | pcDNA5/3FLAG-sirESCO2 T5E | This work |
| pVM11 | pcDNA5/3FLAG-sirESCO2 S244A | This work |
| pVM12 | pcDNA5/3FLAG-sirESCO2 S244E | This work |

\*sir= siRNA resistant

**Table 2:** Antibodies used in this study.

| Antigen | Antibody Source | Number | Dilution | Reference |
| --- | --- | --- | --- | --- |
| acSMC3 | Katsuhiko Shirahige | A7 | 1:1500 | (Nishiyama et al., 2010) |
| β-Tubulin | Development Studies Hybridoma Bank | E7 | 1:4000 |  |
| CDK2 | Cell Signaling | 18048 | 1:1000 |  |
| Cyclin A | Abcam | Ab181591 | 1:2000 |  |
| Cyclin A | BioLegend | 644001 | 1:100 |  |
| ESCO2 | Bethyl Labs | A301-689A | 1:3000 |  |
| Flag | Sigma | F1804-5MG | 1:1000 |  |
| IgG | Jackson | 715035151 | 1:5000 |  |
| MCM4 | Bethyl Labs | A300-193A | 1:3000 |  |
| SMC3 | Lab Made | OMRF160 | 1:3000 | (Song et al., 2012) |
| **Goat a-**<br>**Rabbit (HRP**<br>**conjugated)** | Thermo Fisher<br>Scientific | 31460 | 1:10000 |  |
| **Donkey a-**<br>**Mouse(HRP**<br>**conjugated)** | Jackson<br>ImmunoResearch | 715035151 | 1:5000 |  |

## Acknowledgements

We are grateful to all members of the Rankin lab and the Program in Cell Cycle and Cancer Biology program for many helpful discussions as this project evolved. We are grateful to Michael Davidson for the gift of mEmerald-MCM4-C-19, and Katsuhiko Shirahige for the anti SMC3-Ac antibody. This work was supported by R01GM101250 and R35GM149343 to S.R., and an OMRF predoctoral fellowship to B.R.

---

**Supplemental Figure 1.** Expression of ESCO2 phosphomutant derivatives. Shown is an anti-flag blot of HeLa Flp-In Trex cells engineered to express the indicated flag-tagged ESCO2-derivative transgenes (wild type and phosphomutant). Cells were treated with doxycycline to induce transgene expression, as indicated. SMC3 and tubulin were assessed as loading controls. Cells were treated with ESCO2 siRNA as indicated to deplete endogenous ESCO2 (not shown) in order to score phenotypes attributable to the transgenes. This figure is in support of Figure 1.

