## Supplementary figures and images for "Cell cycle control of cohesion establishment"

### Supplemental Figure 1

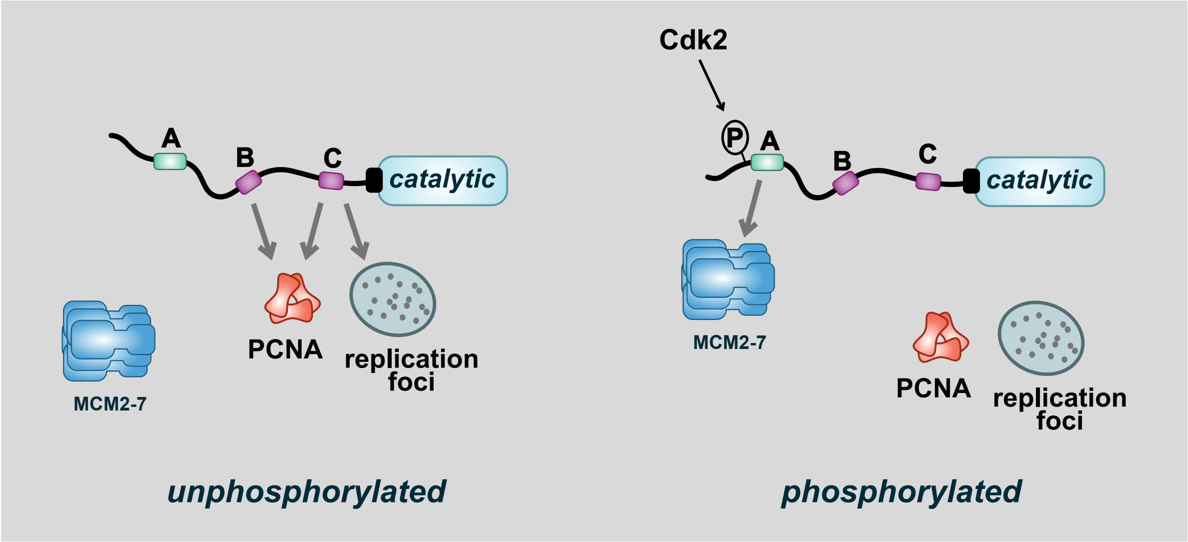
